# *CDM1* is required for meiotic progression and genome integrity during male meiosis in *Arabidopsis*

**DOI:** 10.1101/2025.10.10.681646

**Authors:** Jayeshkumar N Davda, Aswan Nalli, Dipesh K Singh, Keith Frank, Joiselle Fernandes, Imran Siddiqi

**Affiliations:** CSIR Centre for Cellular and Molecular Biology Uppal Road, Hyderabad 500007. India

**Author notes:** Correspondence, Tele, Tele.

**Keywords:** DNA damage inducibility, cell cycle progression, meiotic chromosome organization, microsporogenesis

## Abstract

Microsporogenesis involves the integration of meiosis with the developmental program leading to formation of microspores from meiocytes. The *CALLOSE DEFICIENT MICROSPORE 1* (*CDM1*) gene of *Arabidopsis* has been previously characterized as being required for callose metabolism and its loss results in defects in microspore wall development, degeneration of meiotic products, and male sterility. We report here that CDM1 is required for normal meiosis I progression, meiotic chromosome organization, and maintenance of DNA integrity. CDM1 is part of the ATM dependent DNA damage response in dividing vegetative tissues. The *cdm1* mutant shows defects in meiotic chromosome condensation, arrest at metaphase I, susceptibility to chromosome breakage in rDNA regions, defects in positioning of the products of meiosis, and increased expression of recombination related genes. However, fidelity of chromosome segregation is not affected. Furthermore, the meiotic phenotype of *cdm1* is independent of SPO11 induced meiotic double strand breaks. CDM1 protein displays granular localization in meiosis throughout the cytoplasm that resembles that of mRNP granules. The results show that CDM1 is required for maintenance of proper chromosome organization and DNA integrity in the course of male meiosis. The chromosomal defects in *cdm1* may originate from regions of defective or incomplete replication.

**Highlight:** CDM1 expression is DNA damage inducible in Arabidopsis mitotic cells. CDM1 is required for genome integrity in male meiosis independent of SPO11-induced DNA breaks, suggesting a role in regulation of premeiotic replication repair.

## Introduction

Meiosis is a specialized cell division in which one round of DNA replication is followed by two rounds of cell division leading to halving of chromosome number. During prophase I of meiosis homologous chromosomes undergo pairing and recombination leading to the formation of crossovers. Recombination is initiated by DNA double strand breaks (DSBs) followed by repair of breaks using the homologous chromosome as a template (Murakami and Keeney, 2008). Homologous recombination in meiosis utilizes a core set of proteins that function in repair of DNA damage in somatic cells in addition to context-specific proteins that have specialized functions in meiosis (Krogh and Symington, 2004; Li and Heyer, 2008). For instance, the general eukaryotic strand exchange protein Rad51 is required for DNA repair in somatic cells as well as in meiosis whereas the meiosis-specific strand exchange protein Dmc1 functions with Rad51 and directs repair of Spo11 DNA breaks using the homolog as template instead of the sister chromatid (Brown and Bishop, 2015).

In addition to repairing breaks made by Spo11 in meiosis, DNA repair is also required to prevent accumulation of lesions that lead to loss of chromosomal structure and fragmentation during premeiotic DNA replication. Knockdown in Arabidopsis of the DNA polymerase loading factor CDC45 (Stevens *et al*., 2004) leads to reduced fertility and chromosome fragmentation in meiosis independently of SPO11 breaks whereas vegetative growth of plants under normal conditions is comparable to wild type. Similar SPO11-independent fragmentation phenotypes are observed in the case of *mei1* and *xri1* mutants which are implicated in DNA repair and are part of the DNA damage response in Arabidopsis (Mathilde *et al*., 2003; Dean *et al*., 2009). These lines of evidence suggest that premeiotic DNA replication may be particularly prone to generating lesions which require DNA repair functions for healing and maintenance of chromosomal integrity.

The *CALLOSE DEFECTIVE MICROSPORE 1 (CDM1)* gene of *Arabidopsis* has been characterized primarily with respect to its role in callose metabolism during microspore development in *Arabidopsis* and encodes a protein having a tandem zinc finger (TZF) domain (Lu *et al*., 2014). TZF domain proteins have been extensively studied in mammals and more recently in plants as post-transcriptional regulators of gene expression that bind to RNA and regulate RNA function (Bevilacqua *et al*., 2003; Blackshear *et al*., 2005; Pomeranz *et al*., 2009; Bogamuwa and Jang, 2014; Zhang *et al*., 2019). CDM1 has also been shown to bind RNA and DNA in vitro (Chai *et al*., 2015; O’Malley *et al*., 2016). Here we show that CDM1 has a broader role in meiosis and is required for maintenance of genome integrity as revealed by FISH, and normal progression through male meiosis I. CDM1 is part of the DNA damage response in vegetative tissues undergoing active cell division, however the meiotic phenotype of *cdm1* does not depend on SPO11-induced DNA breaks. CDM1 shows predominantly cytoplasmic localization in meiosis resembling mRNP granules and among genes that are differentially expressed between wild type and the *cdm1* mutant, we observed a signature of recombination and regulated protein turnover via the proteasome. The results suggest that CDM1 plays a role in the prevention or correction of DNA damage or blocks during premeiotic DNA replication.

## Methods

### Plant materials and growth condition

Plants were grown in a plant growth room under long day condition at 22 degrees Celsius as described earlier (Siddiqi *et al*., 2000). The *Arabidopsis* insertion lines having T-DNA insertions in *CDM1*/*AtC3H15*, *SPO11-1-3*, and *ATM* kinase (Supporting Table S1) were obtained from the *Arabidopsis* Biological Resource Center, Ohio. Primers used in this study are listed in Supporting Table S2. Plant transformation was carried out by Agrobacterium (AGL1) mediated in planta transformation (Bechtold and Pelletier, 1998). Transgenic plants were selected on MS-Agar plates containing appropriate antibiotics. For DNA damage treatment, 5-12 day old seedlings were floated in MS media containing 1µg/ml bleomycin for 3 hrs. in the plant growth chamber. Later seedlings were washed thrice with MS and subjected to assay.

### Cloning procedures

The *proCDM1:GFP-GUS-CDM1* construct was made by three way multisite Gateway as per the manual (Thermo Fisher Scientific). The final construct was generated from three pENTR clones, of which the first contained the *CDM1* promoter amplified from genomic DNA with primers CDM1proP4 and CDM1proP1R, the second contained *GFP-GUS* amplified by GFPGUSP1 and GFPGUSP2 primers from pBGWFS7 plasmid (VIB-UGENT CENTRE FOR PLANT SYSTEM BIOLOGY) and the third pENTR clone comprises of the genomic coding and the 3’UTR region of CDM1, amplified by CDM1geneP2R and CDM1 3’UTRP3Rev.

### Cytological procedures

Male meiotic chromosome spreads were prepared according to (Ross *et al*., 1996) with minor modification (Agashe *et al*., 2002). Observations were made on a Zeiss Axioimager imaging microscope, using a Plan Apochromat 63 × oil immersion objective. The male meiotic chromosome spread slides were used for the FISH analysis as described earlier (Singh *et al*., 2013). Centromere and 5S rDNA probes were synthesized using PCR amplification as described earlier (Singh *et al*., 2013). For 45S rDNA probe synthesis, BAC clone f1b23 was digested with Nde1 restriction enzyme to release approximately 10Kb fragment containing 18s, 5.8s, 25s and Internal transcribed spacers. This fragment was used as template for the probe synthesis by nick translation as described earlier (Singh *et al*., 2013).

Immunostaining procedures were followed as described previously (Singh *et al*., 2013) with minor modification. In the case of tubulin immunostaining, inflorescence was fixed in methanol: acetone (3:1) instead of paraformaldehyde. The procedure of immunostaining was carried out using an α-tubulin antibody (Sigma-T5168,1:200) and GFP antibody (ABI-AB6556, 1:200), and all the secondary antibodies were used at 1:200 dilutions. Slides were then mounted in 1 µg/ml DAPI in Vectashield (Vector labs). Slides were observed using a Plan Apochromat 63 × oil immersion objective in Zeiss Axioimager with Apotome imaging microscope for GFP immunostaining and Leica SP8 confocal microscope for the α-tubulin immunostaining. Direct GFP fluorescence was observed in the root of *proCDM1:GFP-GUS-CDM1* complemented plants treated with bleomycin using a 20X objective in a Zeiss-LSM880 confocal microscope. Tissues from *proCDM1:GFP-GUS-CDM1* complemented plants were stained for GUS activity as described earlier (Siddiqi *et al*., 2000). Images were edited with Adobe Photoshop CS3.

### RNA isolation and qRT-PCR

Total RNA was isolated using a plant RNeasy mini kit (Qiagen) following the manufacturer’s protocol. RNA was converted to cDNA by oligo dT primer with Superscript-III enzyme (Invitrogen) as per the manufacturer’s protocol. qRT-PCR was performed in the ABI Prism 7900 HT Sequence Detection System (Applied Biosystems). Cycling parameters consisted of 2 minutes’ incubation at 50°C, 10 minutes at 95°C, and 40 cycles of 95°C for 15 seconds, 60°C for 30 seconds, and 72°C for 30 seconds. GAPC or PDF2 (Czechowski *et al*., 2005; Lilly *et al*., 2011) gene was used as internal normalization standard. Primers used in the study are listed in Supporting Information Table S2.

### Global transcription profiling using microarray

100 ng of total RNA isolated from wild type and *cdm1* meiotic anthers was subjected to analysis on ATH1 microarrays as per the technical manual (Affymetrix). The experiment was repeated with 3 different biological replicates. Data generated after array scan was analyzed using the Expression Console of Affymetrix Gene Chip Operating Software (GCOS). All the CEL (cell intensity) files generated by Expression Console were then loaded into the R Bioconductor “affy” package for microarray analysis (Gautier *et al*., 2004). Probe sets that showed >1.5-fold difference at an FDR corrected p-value < 0.05 were taken as differentially expressed and used for comparative gene expression analysis. Gene enrichment analysis of differentially expressed genes was carried out using the STRING online tool (Szklarczyk *et al*., 2019).

## Results

### CDM1 is expressed in mitotically dividing tissue in response to DNA double strand breaks

Screening of gene expression databases pointed to CDM1 being induced in response to radiomimetic drug treatment that causes DNA double strand breaks (DSBs) (Winter *et al*., 2007; Klepikova *et al*., 2016). We therefore constructed a *proCDM1:GFP-GUS:CDM1* reporter and generated nine *cdm1 proCDM1:GFP-GUS:CDM1* plants in which fertility was restored by the transgene (Supporting Information Figure S1). We then analyzed two complementing lines for reporter expression by assaying GUS activity in parts of the plants and RNA levels in seedlings treated with the radiomimetic drug bleomycin. We found that *CDM1* expression was induced 9-fold in response to bleomycin treatment and this induction was entirely dependent upon the ATAXIA-TELANGECTASIA-MUTATED (ATM) kinase, a master transducer of the DNA DSB damage response in animals and plants (Culligan *et al*., 2006; Tichý *et al*., 2010) (Fig. 1). GUS reporter expression in bleomycin treated seedlings was restricted to root and shoot tips and the basal portion of young leaves (Fig. 1f, g). We did not observe GUS expression in root and shoot tips in untreated seedlings. In agreement with the observations of Lu *et al*., 2014, CDM1 expression was detected in young buds in inflorescences and specifically in anthers. CDM1 expression is therefore part of the ATM-dependent DNA damage response in regions undergoing cell division in vegetative tissue.

**Figure 1.**
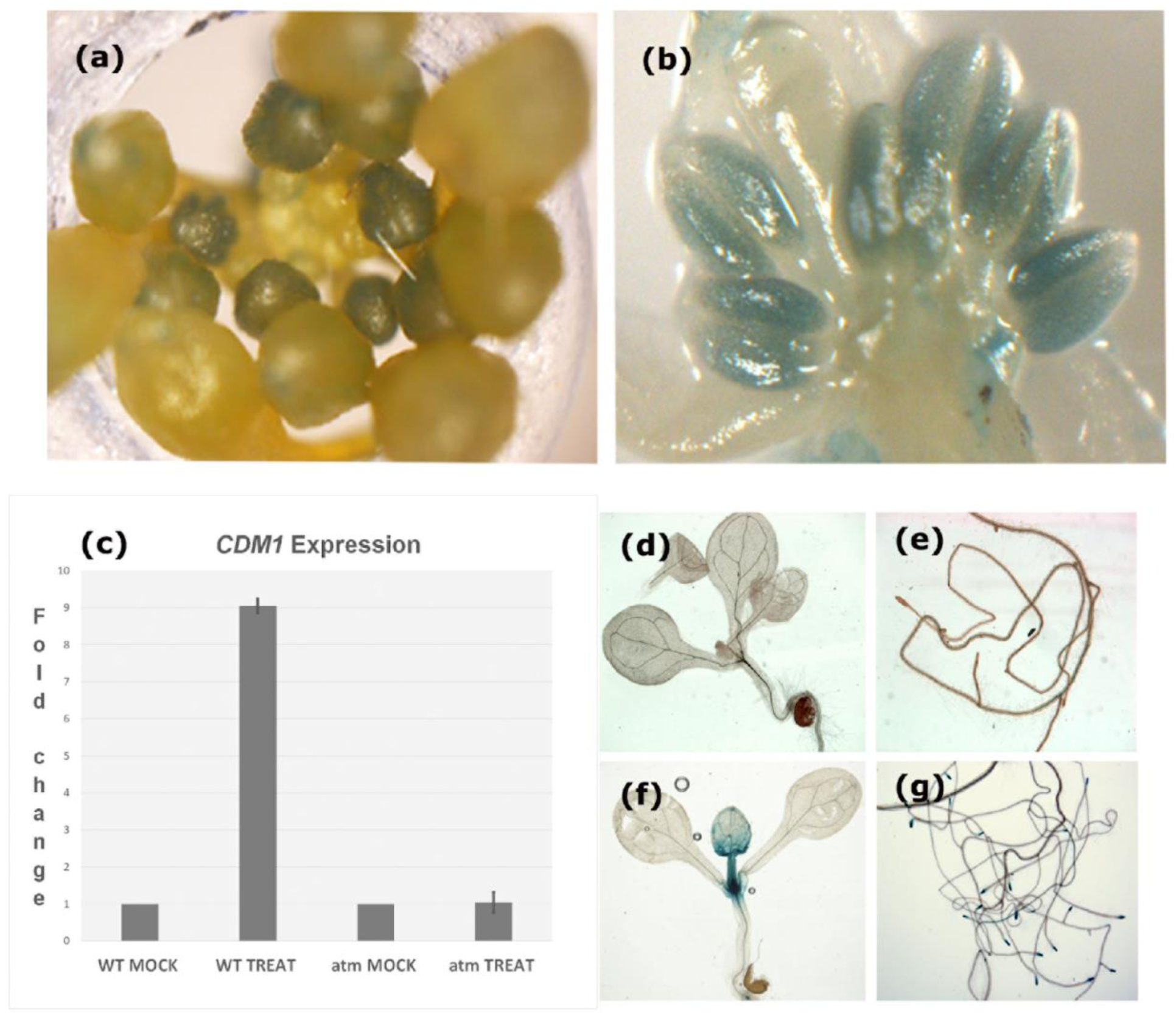
*CDM1* expression during development in young buds and DNA damage induced expression in somatic tissue. (a, b) GUS staining for CDM1 expression in whole inflorescence (a) and anthers of young buds (b). (c) Semi quantitative real-time RT-PCR expression. Fold change is plotted for wildtype and *atm* mutant seedlings treated with bleomycin in comparison to respective mock. (d-g) GUS staining of *_PRO_CDM1:GFP-GUS-CDM1* complemented line, mock shoot (d), mock root (e), bleomycin treated shoot (f), bleomycin treated root (g).

### CDM1 is required for genome integrity, cell cycle progression, and cytokinesis in male meiosis

As CDM1 is part of the response to DNA DSBs in somatic tissue, we examined meiotic chromosome organization in the *cdm1* mutant by performing chromosome spreads on wild type and *cdm1* mutant meiotic stage anthers (Fig. 2). Early and mid-prophase stages of meiosis appeared normal in *cdm1* up to pachytene and chromosomes underwent synapsis and bivalent formation (Supporting Information Figure S2). At diplotene, chromosomes in the *cdm1* mutant showed condensation defects, wherein gaps of uncondensed regions appeared as a thin thread between two condensed regions (Fig. 2). Nonhomologous chromosomes appeared connected by thin threads (Fig. 2e,i) and cases of a separated arm region without visible connection to a whole chromosome were observed (Fig. 2i). At metaphase I the five bivalents had a more extended appearance in *cdm1* in comparison to wild type (Fig. 2f) which further increased at anaphase I (Fig 2g), and at metaphase II the chromosomes appeared as an undercondensed entangled mass. (Fig. 2h). Notwithstanding these condensation defects, chromosome segregation at both meiosis I and meiosis II appeared balanced as Fluorescence in situ hybridization (FISH) using a centromere repeat probe revealed five centromere signals in each of the meiotic products at the end of anaphase I and II (9/9 meiocytes). However, we observed supernumerary DAPI signals in the meiotic products in *cdm1* (Fig 2m-o; 8/9 cases) but not wild type (Fig 2j-l; 0/6 meiocytes). We interpret these signals as arising from acentric fragments either separate from or else connected to centric arms through DAPI invisible undercondensed regions.

**Figure 2.**
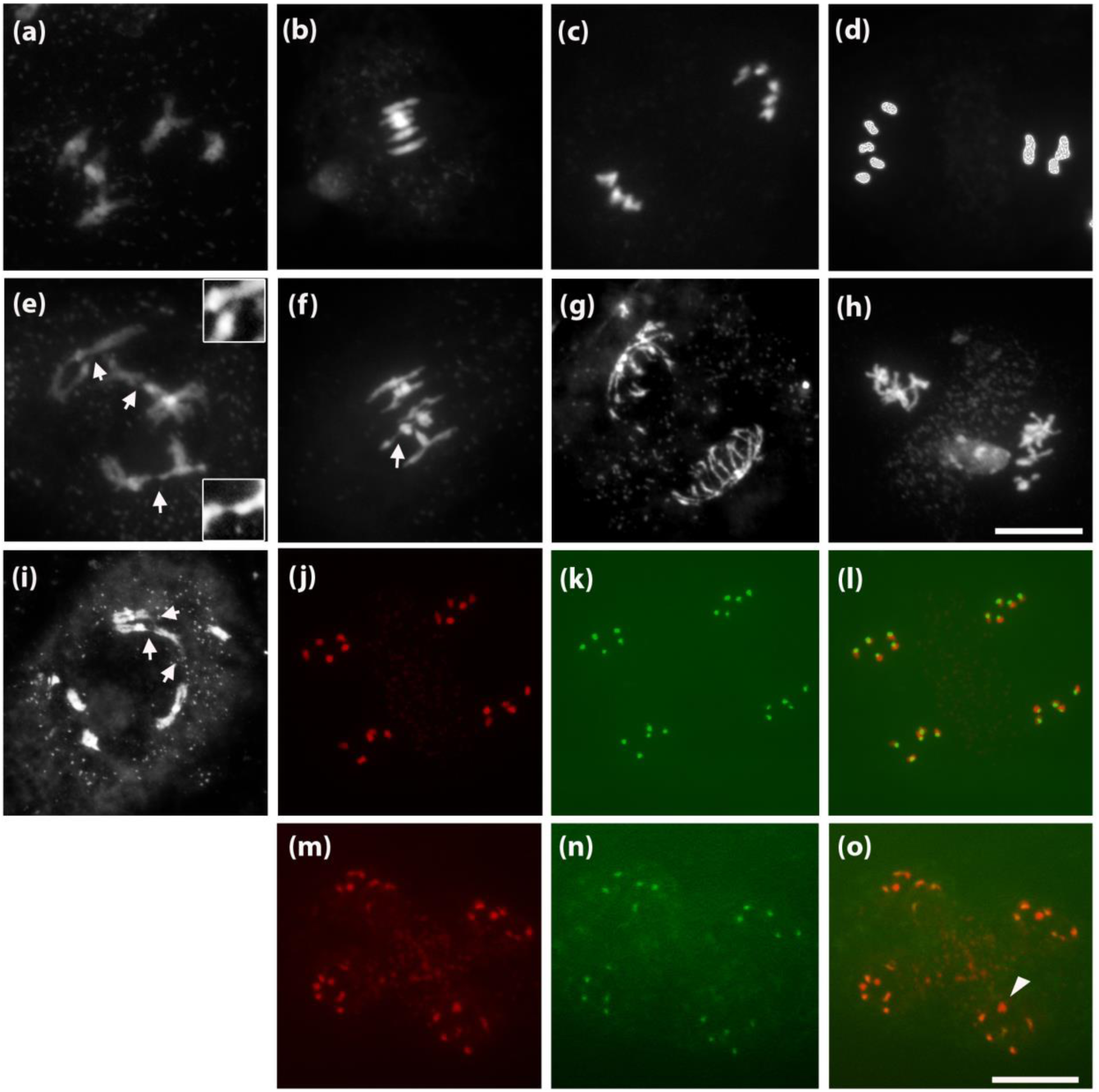
Male meiosis defects in *cdm1* mutant. (a-i) Chromosome spreads of male meiocytes of wild type (a-d) and *cdm1* mutant (e-i). (a,e,i) diplotene, (b,f) metaphase I, (c,g) anaphase I, (d,h) metaphase II. Arrows indicate undercondensed chromosomal region and interchromosomal bridge in *cdm1*. (j-o) Centromeric FISH of telophase II (j,m) DAPI, (k,n) centromere, (l,o) merge. (j-l) Wild type, (m-o) *cdm1*. Arrowhead indicates acentric fragment. Scale Bar 10 µm.

In order to determine if defects in chromosome organization were associated with defective meiotic progression, we quantified meiotic stages in *cdm1* mutant and wild type meiocytes. We observed a 4-fold increase in the proportion of metaphase I stage and a 2.5-fold increase in the proportion of interkinesis stage in *cdm1* indicating that CDM1 is required for progression through meiosis I and *cdm1* mutant meiocytes show arrest at metaphase I (Fig. 3).

**Figure 3.**
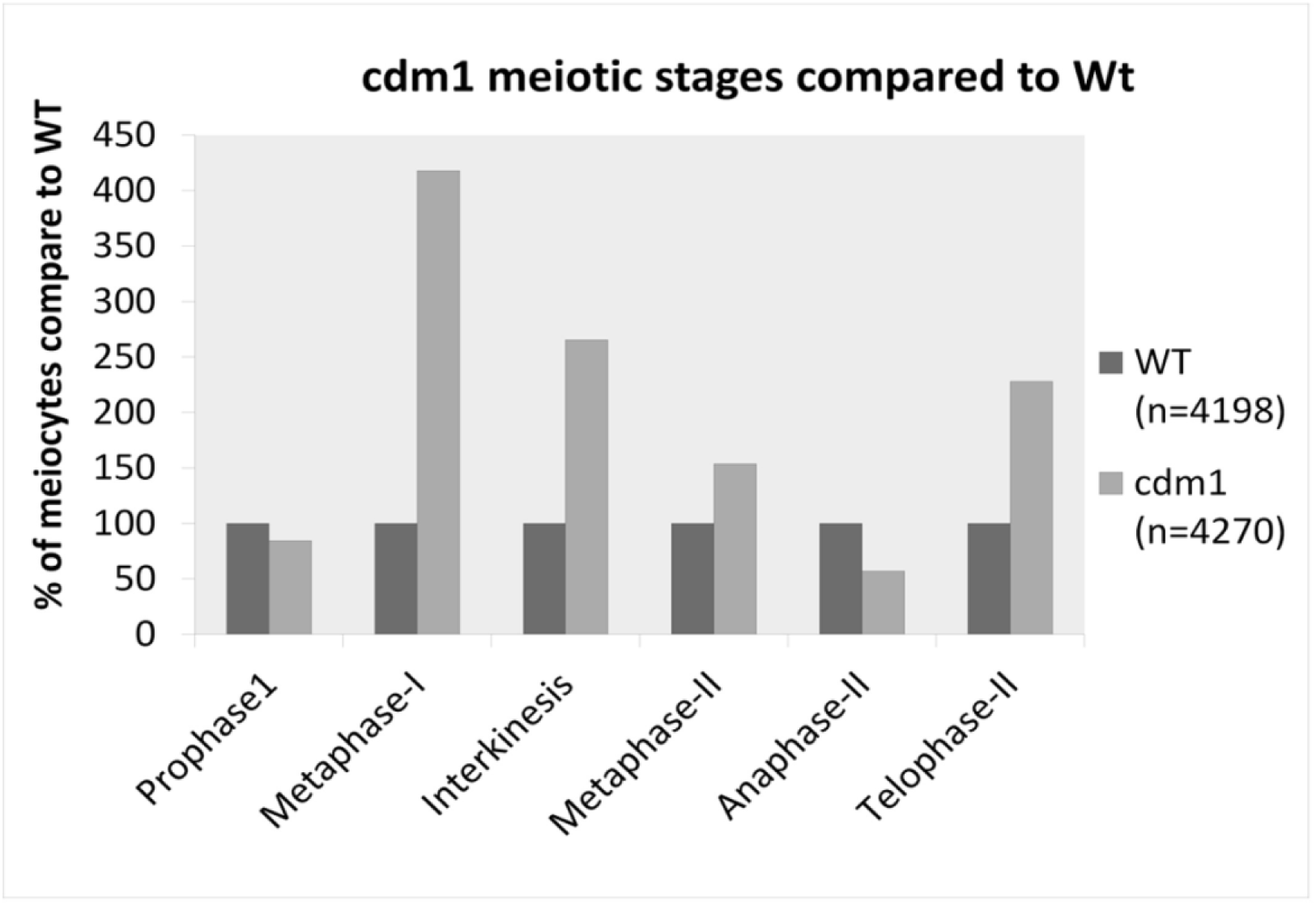
*cdm1* shows metaphase I arrest. Relative roportion of meiotic stages based on counts of chromosome spreads of *cdm1* compared to wild type. For each stage the proportion for wild type was taken as 100%.

Male meiotic cytokinesis in *Arabidopsis* is of the simultaneous type wherein cytokinesis takes place following two rounds of chromosome segregation when four haploid nuclei are located in a tetrad configuration positioned by a radial microtubule array (RMA). The RMA demarcates the cytoplasmic domain of each nucleus (De Storme and Geelen, 2013). Analysis of meiotic products in *cdm1* revealed defects in cytokinesis and irregular cell wall formation (Fig. 4). Meiotic products ranged from tetrads containing one nucleus per cell to all four nuclei being encapsulated in a single cell (Fig. 4f,j) whereas wild type gave only tetrads, pointing to a possible defect in positioning of the nuclei. We, therefore, examined RMA formation in the *cdm1* mutant by staining for tubulin. RMA formation was found to be defective and irregular leading to failure in separation of nuclei at the tetrad stage which would account for encapsulation of more than one nucleus in a single microspore. In some cases, we observed a continuous array extending between nuclei in the case of *cdm1* whereas in wild type each nucleus is associated with its own RMA and there is a gap in the RMAs between adjacent nuclei which marks the zone of cytokinesis. The results described above demonstrate that loss of CDM1 causes a range of defects in meiosis including progression through meiosis I, chromosome condensation and integrity, cell wall biosynthesis, positioning of nuclei after meiosis II, and cytokinesis.

**Figure 4.**
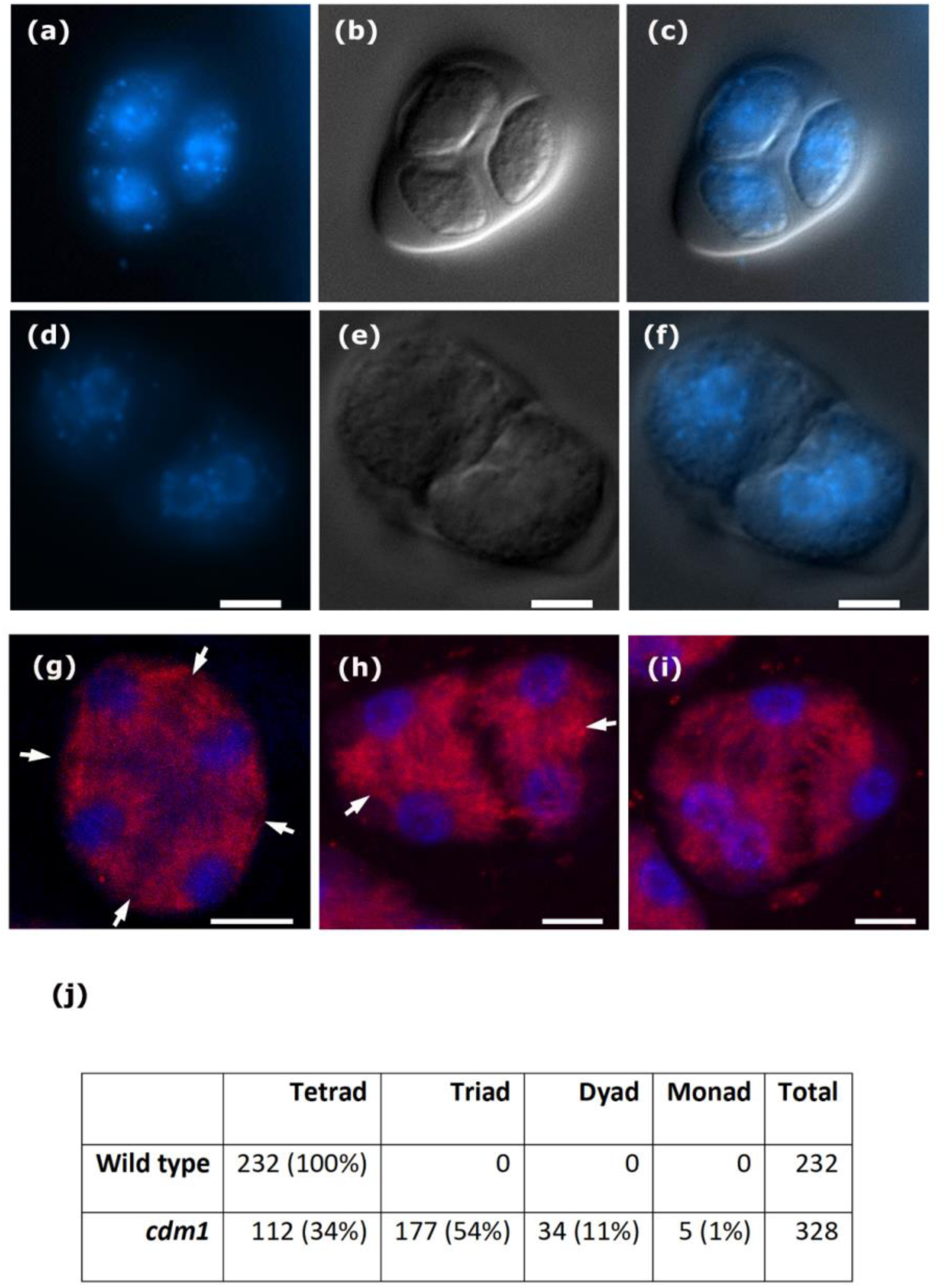
Cytokinesis and RMA formation in wild type and *cdm1* mutant. Cytokinesis in wild type (a-c) and in *cdm1* mutant (d-f). Scale bar 10 µM. RMA formation in wild type (g) and *cdm1* mutant (h and i). Arrowheads indicate difference in discontinuity between wild type and *cdm1*. Scale bar 5 µM. (j) Frequency of meiotic product types in *cdm1* mutant in comparison to wildtype

### The meiotic requirement for CDM1 is not dependent on SPO11-1

The meiotic chromosomal abnormalities in *cdm1* together with the finding that CDM1 is part of the DNA damage response in mitotically dividing cells could be explained if CDM1 plays a role in the repair of DNA double strand breaks made by SPO11 in meiosis (Hartung, 2000; Stacey *et al*., 2006). Meiotic chromosome spreads of *cdm1 spo11-1* meiocytes revealed that the condensation defects observed in *cdm1* were also present in the double mutant indicating that these defects did not arise from failure to repair double strand breaks made by SPO11-1 in meiosis (Fig. 5). Chromosome entanglement and extended appearance in the double mutant at meiosis II was comparable to the *cdm1* single mutant. We examined CDM1 expression in *spo11-1* and *atm* meiotic stage buds and observed that CDM1 expression in male meiosis did not require ATM kinase unlike in mitosis, or DNA double strand breaks made by SPO11-1 (Fig. 4m). CDM1 therefore, has a role in meiotic chromosome development that is not dependent on double strand breaks made by SPO11.

**Figure 5.**
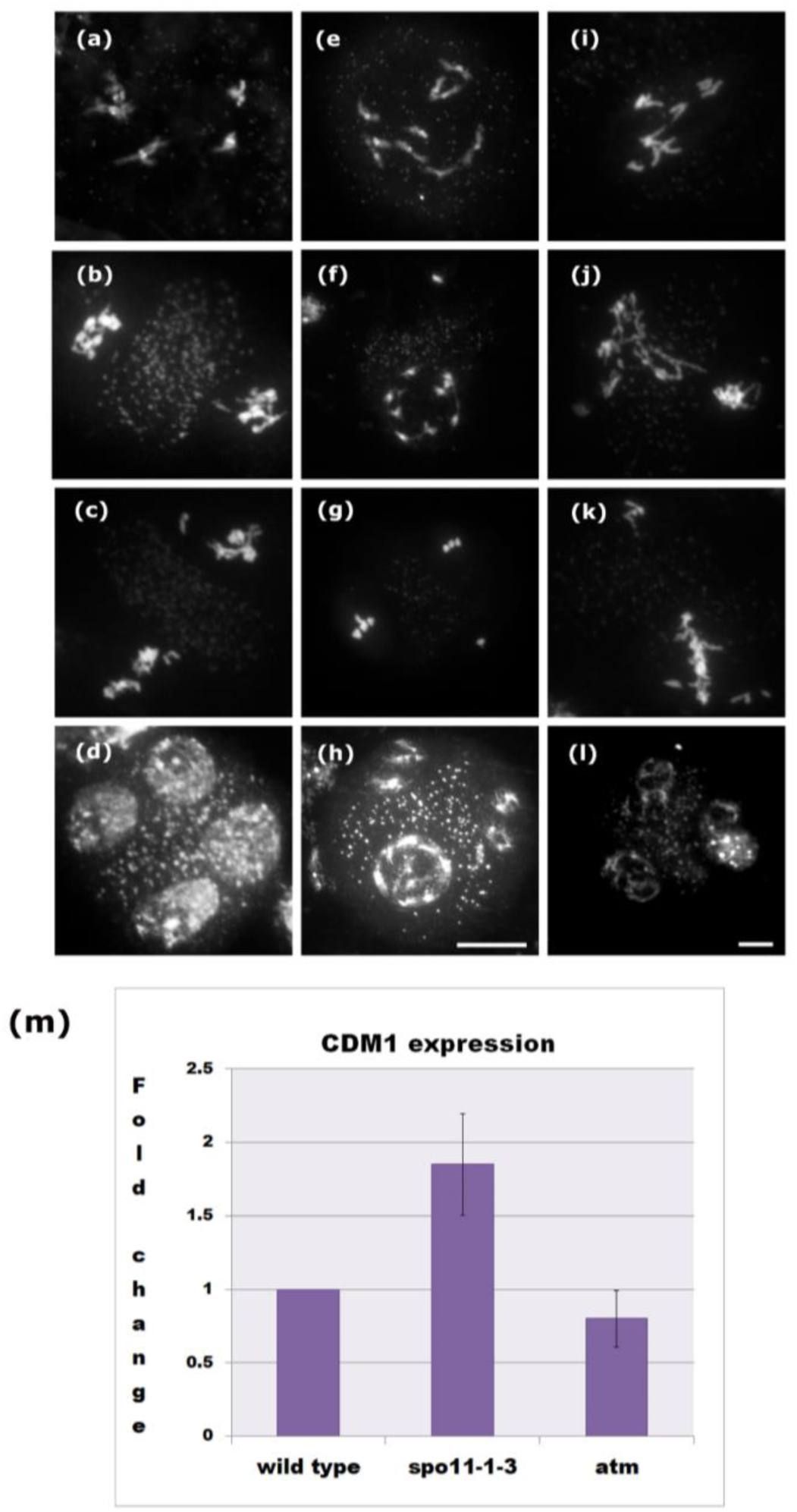
*cdm1* meiotic phenotype and *CDM1* expression is independent of Spo11 mediated DNA DSBs. (a-l) Male meiotic chromosome spreads of of *cdm1*, *spo11*, and *spo11 cdm1* mutants. *cdm1* (a-d), *spo11* (e-h), *spo11 cdm1* (i-l). Diplotene (a, e, and i), Telophase I stage (b, f, and j), metaphase-II (c, g, and k), and telophase II (d, h, and l). Scale bar 10 µm. (m) qRT-PCR for *CDM1* expression in *spo11* and *atm* mutants in comparison to wild type.

### *cdm1* shows defects in rDNA condensation and integrity

A *spo11-1* independent chromosomal phenotype and arrest at metaphase I in *cdm1* suggests the possibility of defects in premeiotic S phase. rDNA sequences are regions of genome instability and sensitive to replication blocks due to the presence of a large number of repeats and conflict between replication and high levels of transcription (Warmerdam and Wolthuis, 2019). We investigated rDNA morphology and integrity in *cdm1* meiosis by rDNA FISH. At diplotene *cdm1* meiocytes showed broad and granular rDNA FISH signals in comparison to wild type suggestive of irregular condensation, and signal was observed even where a DAPI signal was faint or absent (Fig 6b,c). Chromosome 2 and 4 were frequently connected through rDNA region, and these connections persisted through after separation of chromosomes at anaphase I (Fig 6c) suggesting the presence of interchromosomal connections in rDNA regions. These observations point to a requirement for CDM1 in maintenance of rDNA integrity in meiosis.

**Figure 6.**
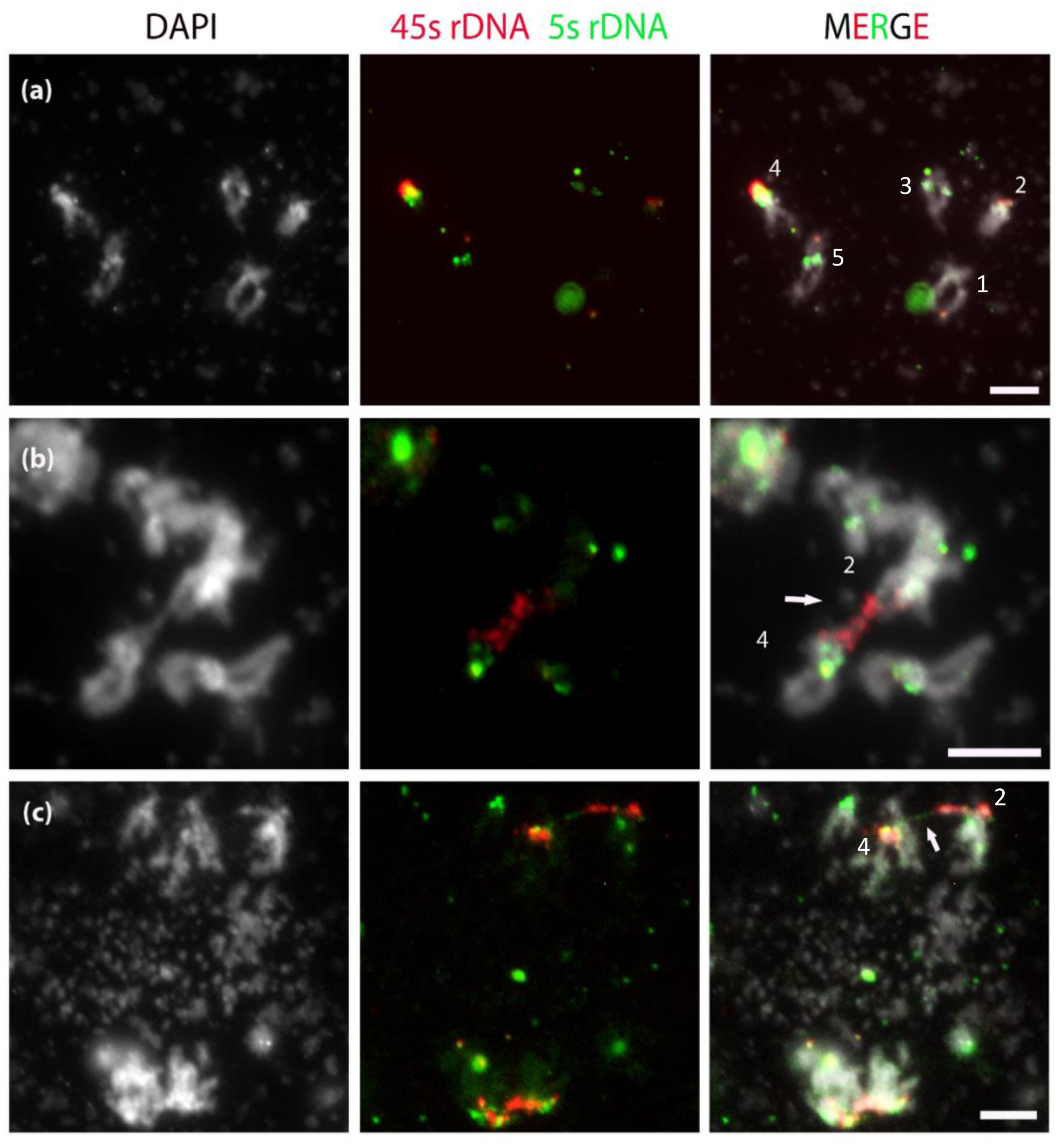
rDNA FISH reveals condensation defects in *cdm1*. (a) Wild type diplotene stage. (b) *cdm1* diplotene. (c) *cdm1* anaphase I. Arrows indicate regions of reduced condensation and chromosomal interconnection through rDNA. Green: 5S rDNA, red: 45S rDNA. (Chr 1: no signal; Chr 2: red; Chr 3: green; Chr 4: green + red: Chr 5: green).

### CDM1 localizes to the nucleus in bleomycin treated roots and to the cytoplasm in male meiocytes

Based on the observed differences in regulation of *CDM1* between mitotic and meiotic cells it was of interest to determine the localization of CDM1 in mitotic vs meiotic cells. We examined the cellular location of the GFP reporter in somatic and meiotic tissue in complemented *proCDM1:GFP-GUS:CDM1* plants (Fig. 7). In the case of meiocytes, indirect immunofluorescence on anther squash was performed as direct GFP fluorescence could not be detected in meiocytes. A prominent GFP signal was detected in the meiotic cells. Localization of the GFP signal in meiocytes was primarily cytoplasmic, extending throughout the cytoplasm, and had a granular appearance. A lower signal was observed in the nucleus with exclusion from the nucleolus. In the case of bleomycin treated seedlings, we were able to directly detect GFP fluorescence in root tips and observed that the signal was mainly in the nucleus where it was concentrated in a central region that likely corresponds to the nucleolus and represents nucleolar sequestration of GFP-GUS:CDM1 (Fig. 5i). CDM1 has been reported to bind both RNA and DNA (Chai *et al*., 2015; O’Malley *et al*., 2016) through its TZF domain and localizes as cytoplasmic foci in transient expression studies in protoplasts (Pomeranz *et al*., 2010). CDM1 binds to a DNA motif which was identified as part of a genomewide analysis of motifs recognized by *Arabidopsis* DNA binding proteins ((O’Malley *et al*., 2016); Supporting Information Fig. S4) and we performed a gene ontology analysis on the occurrence of this motif in the *Arabidopsis* genome using GOMo (Buske *et al*., 2010). The genes identified had a significant enrichment for the GO term “DNA replication initiation” and “DNA unwinding involved in replication” (p = 1.59e-06 and p = 4.98e-05 respectively; Supporting Information Fig. S5) which points to the possibility of CDM1 controlling expression of genes involved in DNA replication.

**Figure 7.**
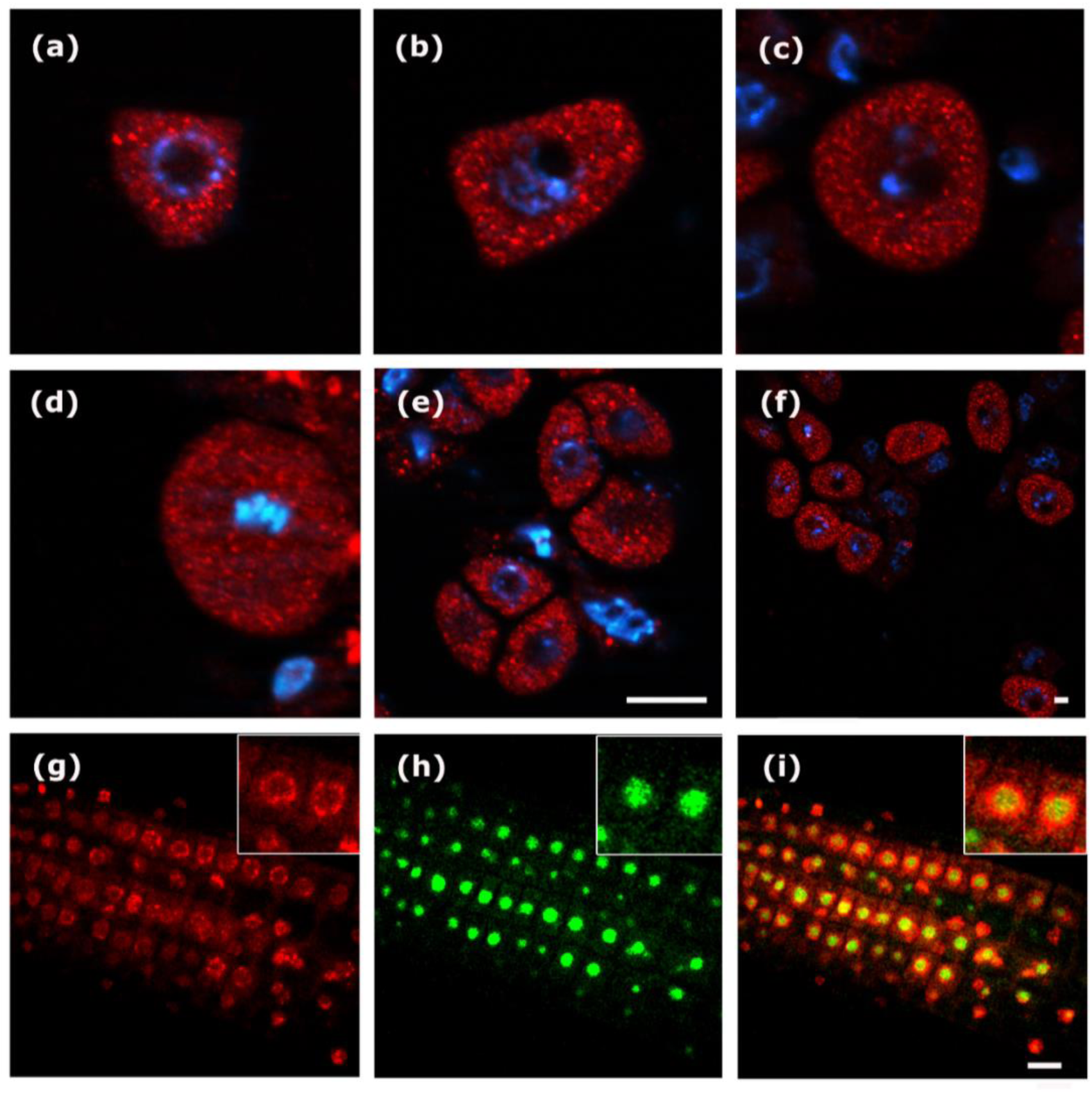
CDM1 localization in meiotic and root tissue. (a-f) Immunostaining against GFP, (red = CDM1, and blue = DAPI), showing localization of CDM1 in different stages of male meiosis. (a) early meiocyte, (b, c) prophase, (d) metaphase I, (e) tetrad, (f) meiocytes staining in comparison to somatic cells. (g-h). GFP fluorescence observed in the root of *_PRO_CDM1:GFP-GUS-CDM1* complemented line upon induction by bleomycin (red = DAPI and green = GFP). Inset photo shows magnified portion of the root cells. Scale Bar 10 µm.

### Expression profiling of *cdm1* reveals signatures of meiotic recombination and controlled protein turnover

In order to examine the changes in gene expression in *cdm1,* we carried out expression profiling using ATH1 microarrays on meiotic stage anthers from *cdm1* and wild type and found 552 genes to be downregulated and 109 to be upregulated in *cdm1* (Fig. 8; fold change > 1.5; p_corr_ < 0.05). We compared the differentially expressed genes we detected to the study by Lu *et al*., (2014) which used RNA from young buds for expression profiling of *cdm1*. We found that 51% of downregulated genes were common, however among upregulated genes there was only 2.6% commonality (Supporting Information Fig. S3). We performed gene ontology (GO) analysis using STRING (Szklarczyk *et al*., 2019) on the differentially expressed genes (Supporting Information Table S3) which revealed strong enrichment among upregulated genes for the biological process GO terms “meiosis I” (FDR = 4.40E-05) and “DNA recombination” (FDR = 8.42E-05). Upregulated genes corresponding to these GO terms included *ASY1*, *DMC1*, *SDS*, *SHOC1*, *AtFANCD2*, *AtMHF1*, *FIGL1*, and *RPA2* all of which play a role in meiotic recombination and repair (Mercier *et al*., 2015). Among down-regulated genes, there was strong enrichment for the biological process GO term “cellular protein modification process” (FDR = 5.68E-05). These included 38 F-box genes (Interpro ID IPR001810; FDR = 6.78E-07). Expression was validated by qRT-PCR for a subset of upregulated and downregulated genes (Fig. 8). To test whether differences in gene expression in *cdm1* were an indirect consequence of progression defects resulting in differences in the representation of meiotic stages, we compared differentially expressed genes in *cdm1* with those in the *mmd1/duet* mutant which has also been shown to cause progression defects in male meiosis and an accumulation of metaphase I stages (Reddy *et al*., 2003; Andreuzza *et al*., 2015; Wang *et al*., 2016) This comparison did not reveal significant overlap suggesting that differentially expressed genes in *cdm1* were unlikely to be due to an indirect consequence of delayed meiotic progression (Supporting Information Fig. S4). The changes in gene expression that are observed in *cdm1* are therefore consistent with defects being at the DNA level (GO term “DNA recombination”) and in cellular transitions (GO term “cellular protein modification process”).

**Figure 8.**
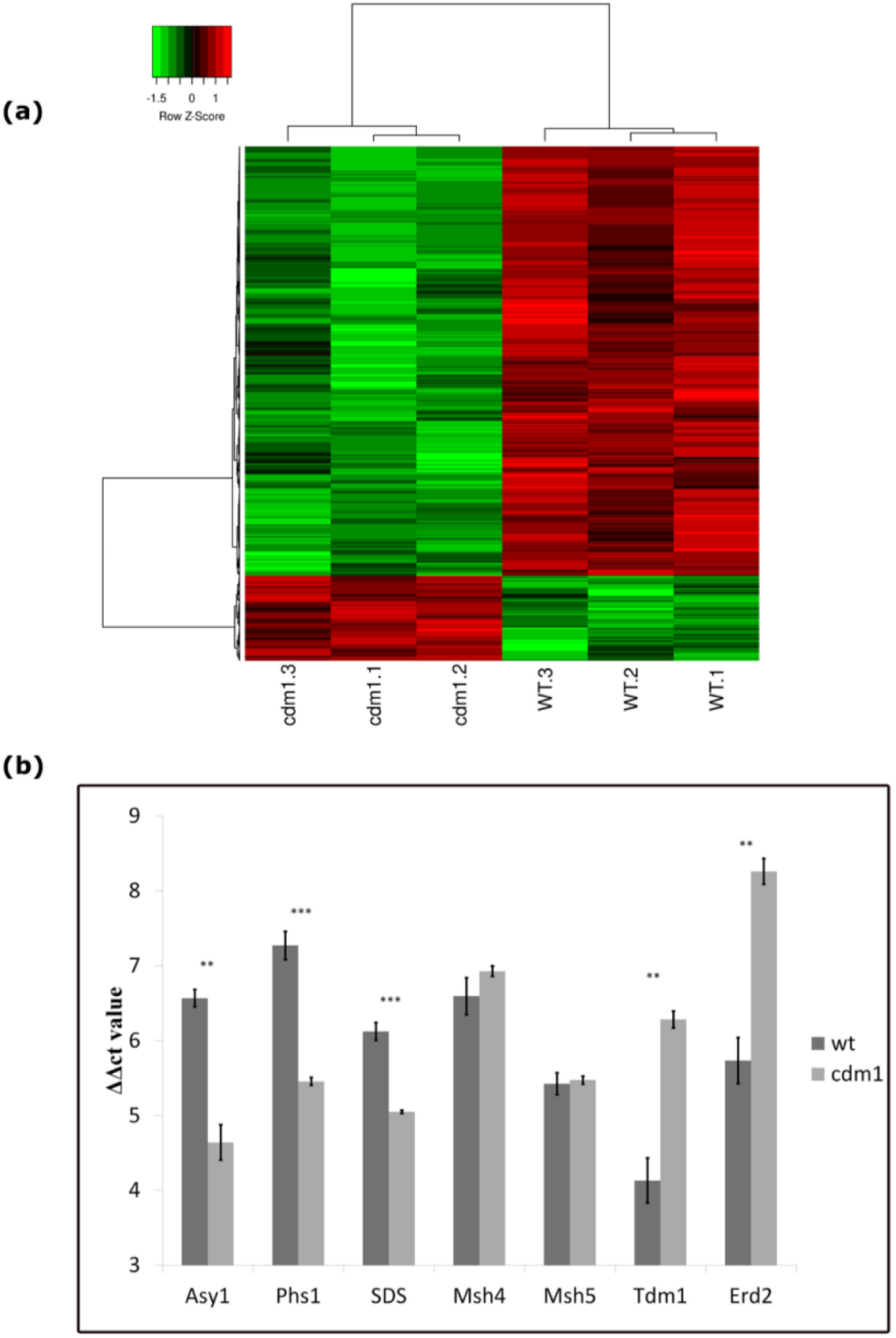
Differentially expressed genes (DEG) in *cdm1* mutant compared to wild type. (a) Heat Map of differentially regulated genes. (b) qRT-PCR validation of selected differentially regulated meiosis specific recombination related genes. *MSH4* and *MSH5* which did not show significant difference in expression were included as a control.

## Discussion

### CDM1 in the DNA damage response and meiosis

We have shown that *CDM1*, a gene required for callose deposition in male meiosis in *Arabidopsis* (Lu *et al*., 2014) is required for the maintenance of meiotic chromosome organization and DNA integrity in the course of male meiosis. CDM1 is part of the DNA damage response in dividing tissue during vegetative growth. *CDM1* expression in seedlings is strongly induced in root and shoot meristematic regions in response to treatment with radiomimetic drug bleomycin and is dependent upon ATM kinase, a master transducer of the DNA damage response (Culligan *et al*., 2006; Ricaud *et al*., 2007). The *cdm1* mutant showed chromosome condensation defects in meiosis I and II. In yeast, unreplicated DNA has been reported to not bind condensin and fail to undergo condensation (Dulev, 2008) and it is possible that the uncondensed regions in *cdm1* also represent underreplicated DNA. We observed a strong arrest phenotype as reflected in a fourfold increase in the proportion of the metaphase I stage, which suggests activation of a G2-M checkpoint in *cdm1*. The observed signature of regulated protein turnover among downregulated genes in *cdm1* may also reflect failure in proper execution of a chromosomal transition in S or G2 of meiosis I. After anaphase I, chromosomes in *cdm1* decondensed at telophase I, however, chromosomes failed to recondense fully in interkinesis and appeared as a tangled group rather than well separated chromosomes at metaphase II. In diakinesis, sister chromatids are in association across their entire length and chromosomes undergo condensation as a pair of sister chromatids, each pair connected to its homolog in the bivalent via chiasmata, whereas in meiosis II after removal of chromosome arm cohesion in meiosis I, sister chromatid arms undergo condensation apart from each other. The prominent condensation phenotype in *cdm1* at metaphase II may reflect defects in organization of sister chromatids interfering with their individual condensation prior to separation at anaphase II. Such entanglement could arise from catenation, topological linkages, or under-replicated DNA (Hirano, 2015; Martinez-Garcia *et al*., 2018). We observed supernumerary DAPI signal at telophase II, however, absence of extensive DNA fragmentation and balanced chromosome segregation in *cdm1* indicates that any such entanglement does not act as a barrier to chromosome separation. We found upregulation of several recombination and repair related genes in *cdm1* which also points to the likely accumulation of defects at the DNA level. These observations indicate that *CDM1* plays an important role in the maintenance of meiotic chromosome integrity and is required for normal progression through meiosis I.

During meiosis, a complex comprising a heterodimer of SPO11-1 and SPO11-2 in conjunction with accessory proteins generates approximately 230 DNA DSBs throughout the *Arabidopsis* genome (Hartung, 2000; Stacey *et al*., 2006; Vrielynck *et al*., 2016). Out of the total, approximately 10 DSBs are processed as crossovers and generate recombinant chromosomes (Chelysheva *et al*., 2007). Interestingly, we found that CDM1 expression and the *cdm1* mutant phenotype in meiosis are not dependent upon SPO11-1 induced DNA double strand breaks. Disruption of the proteins MEI1, CDC45, or XRI1 which have functions connected with DNA replication has been found to cause chromosome fragmentation in meiosis in *Arabidopsis* which is independent of SPO11 and considered to arise from a failure to prevent accumulation of DNA lesions during premeiotic DNA replication (Mathilde *et al*., 2003; Stevens *et al*., 2004; Dean *et al*., 2009). Likewise, the *cdm1* chromosomal phenotype in meiosis may also originate from the accumulation of defects in premeiotic DNA replication. Two additional lines of argument point to the possibility that CDM1 regulates expression of genes involved in DNA replication. Firstly, we found that the DNA motif recognized by CDM1 is enriched at genes that play a role in initiation of DNA replication (Supporting Information Fig. S5; (Buske *et al*., 2010)). TZF proteins are known to act as regulators of RNA stability and it is possible that recruitment through DNA binding can affect the outcome of transcription at these loci. Secondly, CDM1 has been identified as a target of E2F which regulates genes that play a role in the G1-S transition including initiation of DNA replication and in DNA repair (Vandepoele *et al*., 2005).

We observed two different cellular localizations of GFP-GUS:CDM1 in vegetative cells and meiosis. In bleomycin treated seedlings localization of GFP-GUS:CDM1 was mainly nuclear with strong staining in a central region of the nucleus in what likely represents the nucleolus. Recent evidence shows that the nucleolus is a major site of response to DNA damage and a number of DNA repair proteins show nucleolar localization and are either sequestered there or have functions in the nucleolus in repair of rDNA or ribosome biogenesis and mRNA quality control (Ogawa and Baserga, 2017; Kalinina *et al*., 2018; Korsholm *et al*., 2020). In meiosis, localization of GFP-GUS:CDM1 was cytoplasmic and had a granular pattern resembling that of mRNP granules (Cairo *et al*., 2022) and extended throughout the cytoplasm. The *Arabidopsis* protein AtTZF1 has also been reported to traffic between the nucleus and cytoplasmic foci, and to regulate mRNA stability (Qu *et al*., 2014).

### Are chromosomal defects a primary cause of sterility in *cdm1*?

Previous studies have analyzed the function of CDM1 from the perspective of its role in callose metabolism during microspore development and CDM1 has been proposed to regulate the expression of callose synthase and callase genes involved in microspore development (Lu *et al*., 2014; Chai *et al*., 2015). The male sterile phenotype of *cdm1* has been ascribed directly to the failure in regulation of callose metabolism. The finding of meiotic chromosome condensation and progression defects in *cdm1* that manifest in meiosis I before the cytokinesis and tetrad stages lead us to consider an alternative explanation for the *cdm1* phenotype: that the defect in callose deposition, RMA formation, and cytokinesis is a downstream consequence triggered by a defect in a step of meiotic chromosome development. Reports of mutants affecting both meiotic chromosome organization as well as cytokinesis or RMA formation in microspore development include the *rsw4* allele of *AESP* encoding Arabidopsis Separase which has been shown to affect meiotic chromosome disjunction and RMA formation (Yang *et al*., 2011), and *cdkg1-1* which affects somatic and meiotic recombination (Nibau *et al*., 2020) as well as callose deposition in microspores (Huang *et al*., 2013). CDKG1 associates with splicing factors and controls splicing of CALS5 (Huang *et al*., 2013). Our results together with existing lines of evidence suggest the possibility that the integrity of meiotic chromosome organization and segregation is closely connected to control of RMA and cell wall formation in microspore development. In the light of the overall evidence we speculate that the phenotypes of *cdm1* likely initiate from accumulation of defects in premeiotic DNA replication.

In summary, we have shown that *CDM1*, a gene so far characterized primarily for its role in cell wall formation during microsporogenesis, is part of the DNA damage response in mitotically dividing vegetative cells and is required for normal meiotic chromosome condensation and progression through meiosis I. The analysis shows that CDM1 plays an important role in meiotic chromosome organization and that its action is at a stage distinct from repair of SPO11 induced double strand breaks, possibly going back to pre-meiotic DNA replication. Our findings are consistent with CDM1 playing a role in regulation of gene expression in meiosis starting from premeiotic DNA replication, through meiosis I, to cell wall formation at the end of meiosis.

## Supplementary data

Supporting Information Figure S1: Complementation of *cdm1* mutant with *proCDM1:GFP-GUS:CDM1* construct and restoration of fertility.

Supporting Information Figure S2: Meiotic prophase chromosomes of wild type and *cdm1*. (a-c) Wild type, (d-f) *cdm1*. (a,d) leptotene; (b,e) zygotene; (c,f) pachytene

Supporting Information Figure S3: Comparison of differentially expressed genes in *cdm1* mutant with previously published data of *cdm1* mutant.

Supporting Information Figure S4: comparisons of differentially expressed genes in *cdm1* mutant and *duet*.

Supporting Information Figure S5: Gene ontology analysis of the CDM1 binding motif in promoters of all genes in *Arabidopsis*.

Supporting Information Table S1: T-DNA insertion lines used in the study.

Supporting Information Table S2: Primers used in the study.

Supporting Information Table S3: STRING analysis of differentially expressed genes in *cdm1* mutant compared to wild type (a separate excel file is attached).

Supporting Information Table S4: Gene description of top 10 genes found in GOMo analysis.

## Acknowledgements

We thank Madhavi, Ramesh and Chandan Kumar for the assistance with Microarray experiment. JND thanks Kavya Y and Avinash Kumar Singh for the technical assistance, and Jyotsna Dhawan for comments on the manuscript.

## Author’s contribution

JND, IS: Conceptualization; JND, DKS, KF, JF: Investigation; AN, JND: Resources; JND, IS: Writing; JND, DKS, IS: visualization; IS: supervision; IS: Funding acquisition.

## Data availability Statement

The data supporting the findings of this study are available from the corresponding author, Dr. Imran Siddiqi, upon request.

