## Supplementary Information for "*CDM1* is required for meiotic progression and genome integrity during male meiosis in *Arabidopsis*"

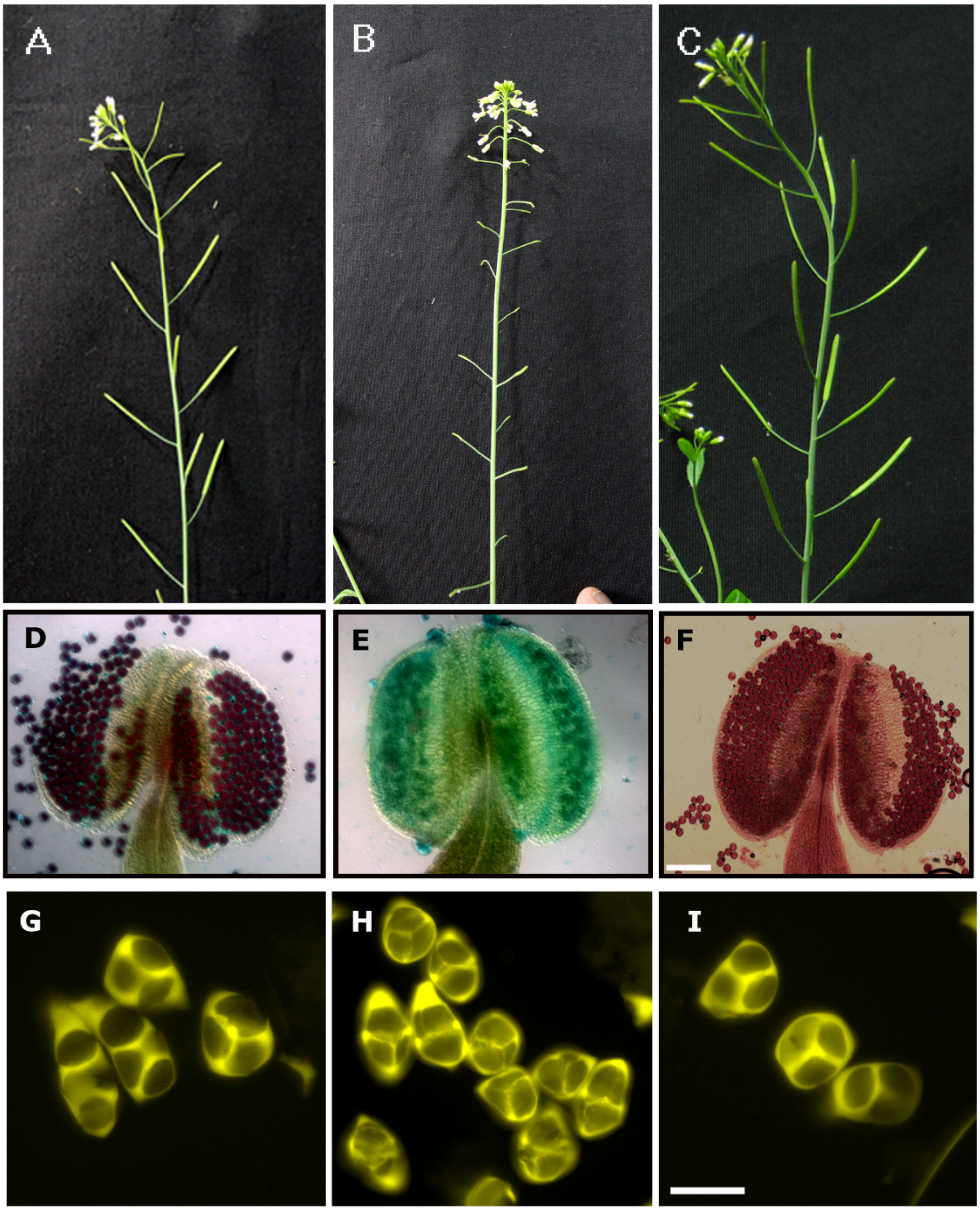


**Supporting Information Figure S1: Complementation of cdm1 mutant with *proCDM1:GFP-GUS:CDM1* construct and restoration of fertility**. (A, D, and G) wildtype, (B, E, and H) *cdm1* mutant sterile plant, (C, F, and I) complementation restores fertility, (A-C) inflorescence and siliques, (D-F) Alexander staining for pollen viability, (G-I) aniline blue staining for callose.

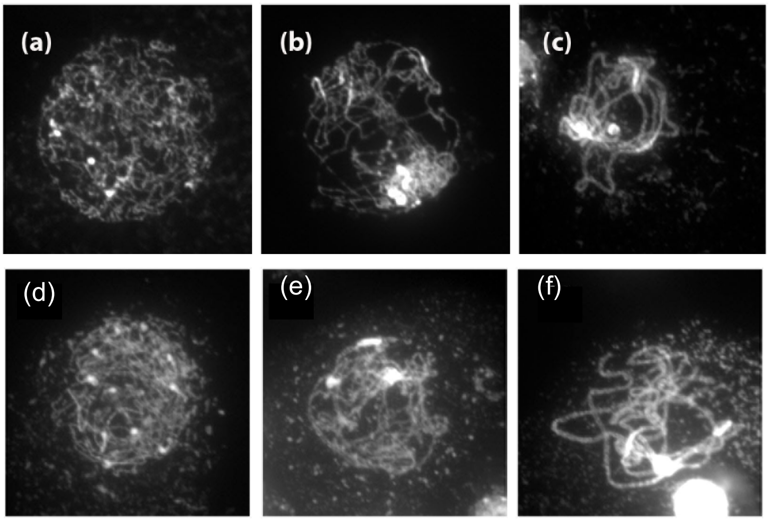


**Supporting Information Figure S2: Meiotic prophase chromosomes of wild type and *cdm1*.** (a-c) Wild type, (d-f) *cdm1*. (a,d) leptotene; (b,e) zygotene; (c,f) pachytene

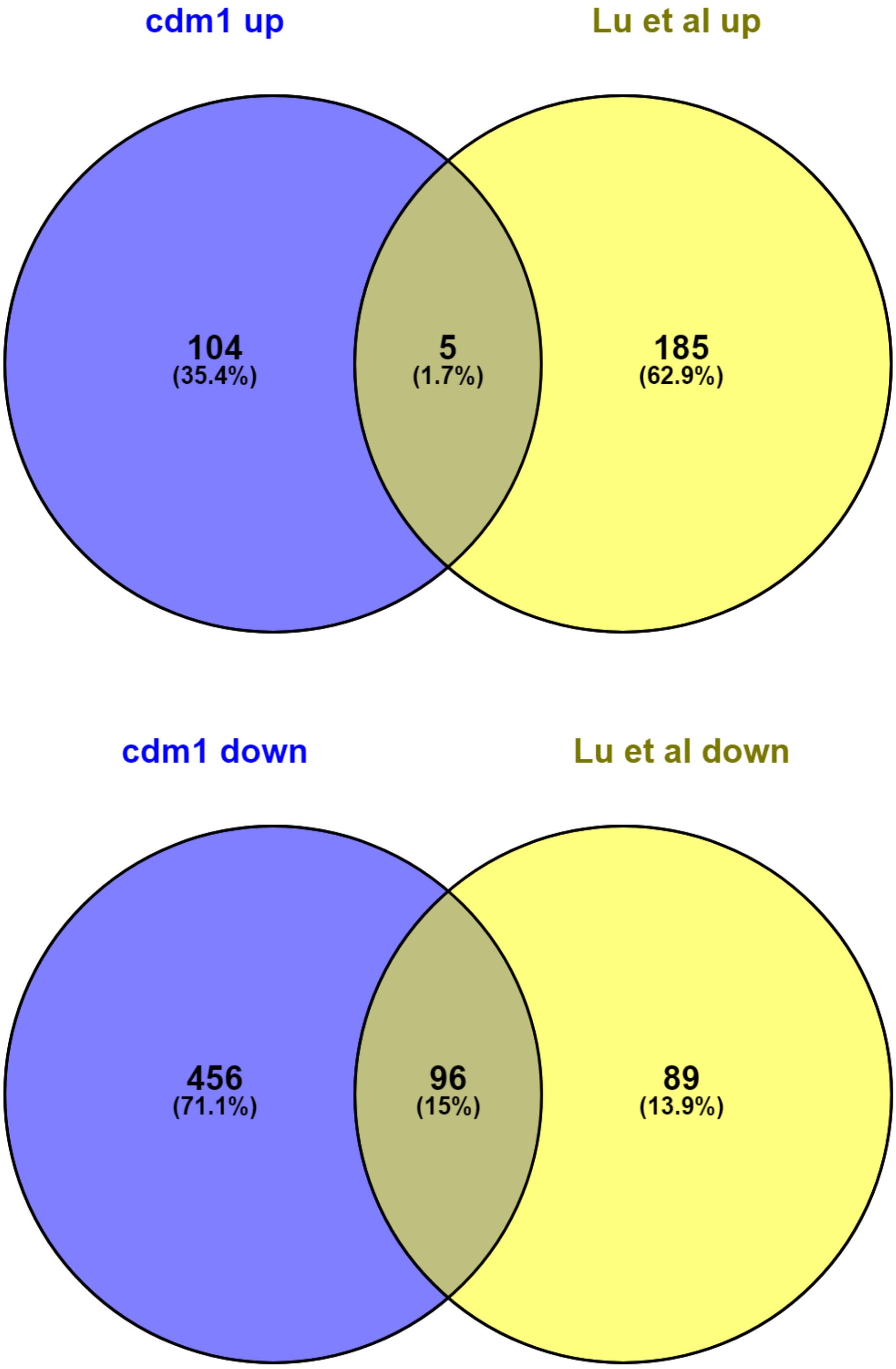


**Supporting Information Figure S3: Comparison of differentially expressed genes in *cdm1* mutant with previously published data of *cdm1* mutant.**

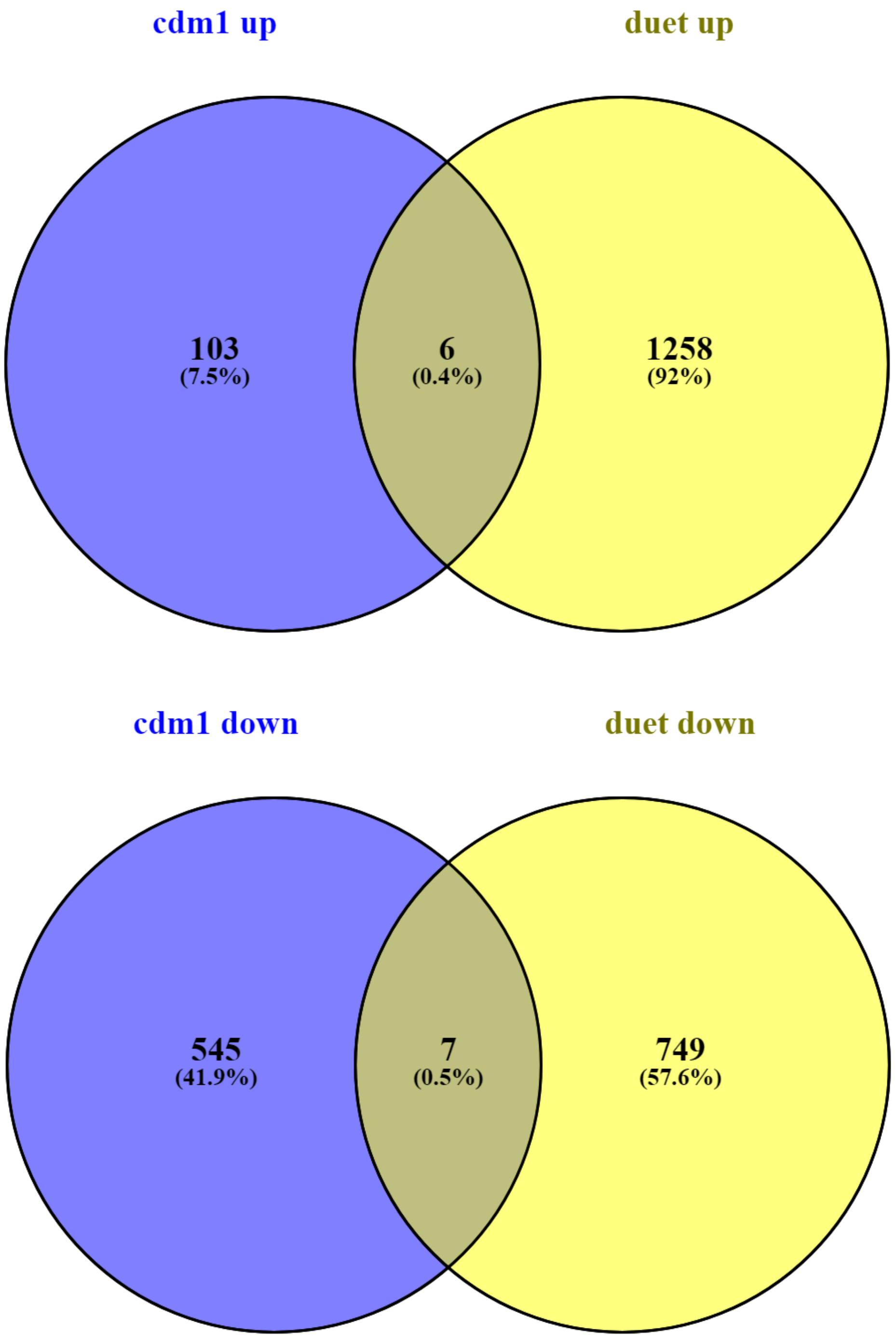


**Supporting Information Figure S4: comparisons of differentially expressed genes in *cdm1* mutant and *duet*.**

### GOMo
Gene Ontology for Motifs

For further information on how to interpret these results please access https://meme-suite.org/meme/doc/gomo-output-format.html.
To get a copy of the MEME software please access https://meme-suite.org.

If you use GOMo in your research, please cite the following paper:
Fabian A. Buske, Mikael Bodén, Denis C. Bauer and Timothy L. Bailey, "Assigning roles to DNA regulatory motifs using comparative genomics", *Bioinformatics*, 26(7), 860-866, 2010. [full text]

INVESTIGATED MOTIFS | PROGRAM INFORMATION | RESULTS IN TSV FORMAT | RESULTS IN XML FORMAT

#### Investigated Motifs

##### Overview

| Motif | Logo | Predictions | Top 5 specific predictions |
|---|---|---|---|
| AT1G68200 | 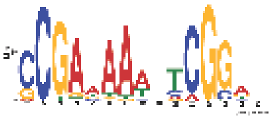 | 4 | CC mitochondrion<br>CC chloroplast<br>BP DNA replication initiation<br>BP DNA unwinding involved in replication |

#### Motif AT1G68200

Top

GO terms are shown in grey if a more specific GO term was also significantly associated with this motif. The most specific GO terms are shown in black.
BP stands for biological process, CC stands for cellular component and MF stands for molecular function.

| GO term | score | *p*-value | *q*-value | Specificity | GO name | Gene ID / Rank (25649 genes in total) |
|---|---|---|---|---|---|---|
| GO:0005739 | 3.187e-03 | 2.652e-07 | 4.996e-04 | ~12% | CC mitochondrion | AT5G52060 (12), AT4G04870 (19), AT2G14095 (57), AT1G55860 (102), AT1G72170 (123), AT4G05136 (168), AT3G20040 (192), AT2G44950 (194), AT2G46540 (216), AT2G45030 (218), ...873 more... |
| GO:0009507 | 4.695e-03 | 2.652e-07 | 4.996e-04 | 20% | CC chloroplast | AT1G16880 (4), AT3G17830 (9), AT2G39140 (17), AT3G59640 (24), AT4G30310 (28), AT1G33780 (30), AT4G04480 (35), AT5G43770 (37), AT2G02980 (41), AT4G31870 (43), ...1957 more... |

| GO term | score | p-value | q-value | Specificity | | GO name | Gene ID / Rank (25649 genes in total) |
|---|---|---|---|---|---|---|---|
| GO:0006270 | 1.239e-02 | 1.591e-06 | 1.998e-03 | ~95% | BP | DNA replication initiation | AT3G09660 (140), AT2G20980 (227), AT5G44635 (474), AT4G02060 (1072), AT1G80190 (1304), AT2G07690 (2315), AT5G46280 (2726), AT5G49010 (3025), AT2G14050 (3839), AT1G44900 (10720), ...1 more... |
| GO:0006268 | 2.988e-02 | 4.985e-05 | 4.696e-02 | 100% | BP | DNA unwinding involved in replication | AT5G44635 (474), AT4G02060 (1072), AT5G63920 (1433), AT2G07690 (2315), AT5G46280 (2726), AT4G31210 (7801), AT2G32000 (10509), AT1G44900 (10720), AT2G16440 (12994) |

**GOMo version:**
5.4.1 (Release date: Sat Aug 21 19:23:23 2021 -0700)

**Input Data:**
go-term-sequence mapping: db/plant_arabidopsis_1000_199.na.csv
scored sequence file: plant_arabidopsis_1000_199.na.cisml
scored sequence file: plant_oryza_sativa_1000_199.na.cisml
scored sequence file: plant_populus_trichocarpa_1000_199.na.cisml
scored sequence file: plant_sorghum_brachupodium_1000_199.na.cisml
scored sequence file: plant_brachypodium_1000_199.na.cisml

**Command line summary:**
This information can also be useful in the event you wish to report a problem with the GOMo software.

Command:

```
gomo --nostatus --verbosity 1 --oc . --t 0.05 --shuffle_scores 1000 --dag db/go.dag --motifs
CDM1_binding_motif.txt db/plant_arabidopsis_1000_199.na.csv
plant_arabidopsis_1000_199.na.cisml plant_oryza_sativa_1000_199.na.cisml
plant_populus_trichocarpa_1000_199.na.cisml plant_sorghum_brachupodium_1000_199.na.cisml
plant_brachypodium_1000_199.na.cisml
```

Seed: 32795697

Significance Threshold: 0.05



**Supporting Information Figure S5: Gene ontology analysis of the CDM1 motif in promoters of all genes in Arabidopsis.**

**Supporting Information Table S1: T-DNA insertion used in the study.**

| **Gene** | **Salk line ID** |
|---|---|
| *AtC3H15/CDM1* | SALK_065040 (Lu *et al.*, 2014) |
| *Spo11-1-3* | SALK_146172 (Sanchez-Moran *et al.*, 2007) |
| *ATM Kinase* | SALK_089805 |

**Supporting Information Table S2: Primer used in the study.**

| Primer | Sequence (5'-3') |
|---|---|
| CDM1LP | TAAATCTGGCATGCAAATATG |
| CDM1RP | TGAATTCACCACATAACCGATG |
| SALK_LB1.3 | ATTTTGCCGATTTCGGAAC |
| ATMLP | CCAAACAAAATCGTTAGCCTG |
| ATMRP | CGAGGGTGTAGCCATATTCAC |
| Spo11-1-3LP | AATCGGTGAGTCAGGTTTCAG |
| Spo11-1-3RP | CCATGGATGAAAGCGATTTAG |
| CDM1RTFWD | TGAATCGCTCTTCGCTTCCA |
| CDM1RTREV | TTCCCTCCTCCTCGCACATA |
| CDM1proP4 | GGGGACAACTTTGTATAGAAAAGTTGTATGGTCCC AAGAAAATGGCAGAGTAACAC |
| CDM1proP1R | GGGGACTGCTTTTTTGTACAAACTTGTCATTTTTCC CGGTGACAATTCTGTTACAC |
| GFPGUSP1 | GGGGACAAGTTTGTACAAAAAAGCAGGCTACATG GTGAGCAAGGGCGAGGAGCTGTT |
| GFPGUSP2 | GGGGACCACTTTGTACAAGAAAGCTGGGTATCATT GTTTGCCTCCCTGCTGC |
| CDM1geneP2R | GGGGACAGCTTTCTTGTACAAAGTGGAAATGGAA AACAAAATCGCGCCGTTTAG |
| CDM1 3'UTRP3Rev | GGGGACAACTTTGTATAATAAAGTTGATGTTGTAA ACCGTGGTAGTCAAACTCTC |
| GAPC-qR2 | CTTGAGTTTGCCTTCGGATT |
| GAPC-qF2 | CCGTTGATGTCTCAGTTGTTG |
| TDM1RTF | TCGTCGGAGAGACGTGATCC |
| TDM1RTR | ATCAACTCGATCTCCAGCGTT |
| SDSRTF | GCCTGCATCGAACACAACAA |
| SDSRTR | TTACTGCCCAAGCAACCAGT |
| MSH4RTF | GCATGGTTGGGGTATCGGAA |
| MSH4RTR | GCCAAAGGTTCTTCTGCTGC |
| MSH5RTF | ATTCAGCTACGAGCAAGCGT |
| MSH5RTR | GACTTCACTGCCCACATCCA |
| ASY1-2RTF | TAGCAAGGGTGAAGGACTGG |

| | |
|---|---|
| ASY1-2RTR | GGTGAATTCGCTCTCTGGTG |
| PHS1RTF | GCTTCCCCGACTCCACTCTA |
| PHS1RTR | ATCAGGTAATCCAGTTACCTCCAA |
| TDM1RTF | TCGTCGGAGAGACGTGATCC |
| TDM1RTR | ATCAACTCGATCTCCAGCGTT |
| SDSRTF | GCCTGCATCGAACACAACAA |
| SDSRTR | TTACTGCCCAAGCAACCAGT |
| MSH4RTF | GCATGGTTGGGGTATCGGAA |
| MSH4RTR | GCCAAAGGTTCTTCTGCTGC |
| ERD2-2RTF | TCGTACGTTGCTTTCACCGA |
| ERD2-2RTR | GGGTTACCTTGAAGGGCCAA |
| PDF2F | TCATTCCGATAGTCGACCAAG |
| PDFR | TTGATTTGCGAAATACCGAAC |

**Supporting Information Table S3: STRING analysis of differentially expressed gene in cdm1 mutant compared to wild type (a separate excel file is attached).**

**Supporting Information Table S4: Gene description of top 10 genes found in GOMo analysis.**

| Locus Identifier | Gene Description | Gene Model Type | Primary Gene Symbol | All Gene Symbols |
|---|---|---|---|---|
| AT4G31210 | DNA topoisomerase, type IA, core;(source:Araport11) | protein_coding | | |
| AT2G32000 | DNA topoisomerase, type IA, core;(source:Araport11) | protein_coding | | |
| AT1G44900 | Encodes MCM2 (MINICHROMOSOME MAINTENANCE 2), a protein essential to embryo development. Overexpression results in altered root meristem function. | protein_coding | MINICHROMOSOME MAINTENANCE 2 (MCM2) | MINICHROMOSOME MAINTENANCE 2 (MCM2)<br><br>(ATMCM2) |
| AT5G46280 | Involved in the replication of mungbean yellow mosaic India virus (MYMIV) DNA through an ex vivo system. | protein_coding | MINICHROMOSOME MAINTENANCE 3 (MCM3) | MINICHROMOSOME MAINTENANCE 3 (MCM3) |
| AT2G16440 | Regulates DNA replication via interaction with BICE1 and MCM7. | protein_coding | MINICHROMOSOME MAINTENANCE 4 (MCM4) | MINICHROMOSOME MAINTENANCE 4 (MCM4) |
| AT2G07690 | Member of the minichromosome maintenance complex, involved in DNA replication | protein_coding | MINICHROMOSOME MAINTENANCE 5 (MCM5) | MINICHROMOSOME MAINTENANCE 5 (MCM5) |

| | | | | |
|---|---|---|---|---|
| | initiation. Abundant in proliferating and endocycling tissues. Localized in the nucleus during G1, S and G2 phases of the cell cycle, and are released into the cytoplasmic compartment during mitosis. Binds chromatin. | | | |
| AT5G44635 | minichromosome maintenance (MCM2/3/5) family protein;(source:Araport11) | protein_coding | MINICHROMOSOME MAINTENANCE 6 (MCM6) | MINICHROMOSOME MAINTENANCE 6 (MCM6) |
| AT3G09660 | Encodes a minichromosome maintenance protein that is involved with RAD51 in a backup pathway that repairs meiotic double strand breaks without giving meiotic crossovers when the major pathway, which relies on DMC1, fails. | protein_coding | MINICHROMOSOME MAINTENANCE 8 (MCM8) | MINICHROMOSOME MAINTENANCE 8 (MCM8)<br><br>(ATMCM8) |
| AT2G14050 | minichromosome maintenance 9;(source:Araport11) | protein_coding | MINICHROMOSOME MAINTENANCE 9 (MCM9) | MINICHROMOSOME MAINTENANCE 9 (MCM9) |
| AT1G80190 | Similar to the PSF1 component of GINS complex, which in other organism was shown to be involved in the initiation of DNA replication. | protein_coding | PARTNER OF SLD FIVE 1 (PSF1) | PARTNER OF SLD FIVE 1 (PSF1) |

| | | | | |
|---|---|---|---|---|
| AT4G02060 | Member of the minichromosome maintenance complex, involved in DNA replication initiation. Abundant in proliferating and endocycling tissues. Localized in the nucleus during G1, S and G2 phases of the cell cycle, and are released into the cytoplasmic compartment during mitosis. Binds chromatin. | protein_coding | PROLIFERA (PRL) | PROLIFERA (PRL)<br><br>(MCM7) |
| AT5G49010 | Similar to the SLD5 component of GINS complex, which in other organism was shown to be involved in the initiation of DNA replication. | protein_coding | SYNTHETIC LETHALITY WITH DPB11-1 5 (SLD5) | SYNTHETIC LETHALITY WITH DPB11-1 5 (SLD5)<br><br>EMBRYO DEFECTIVE 2812 (EMB2812) |
| AT5G63920 | Encodes topoisomerase 3alpha. Suppresses somatic crossovers. Essential for resolution of meiotic recombination intermediates. | protein_coding | TOPOISOMERASE 3ALPHA (TOP3A) | TOPOISOMERASE 3ALPHA (TOP3A)<br><br>(AtTOP3alpha) |
